# Age-Volume Associations in Cerebellar Regions by Sex and Reproductive Stage

**DOI:** 10.1101/2021.11.04.467177

**Authors:** Tracey H. Hicks, Hannah K. Ballard, Huiyan Sang, Jessica A. Bernard

## Abstract

**Introduction:** The cerebellum has established associations with motor function and a well-recognized role in cognition. In advanced age, cognitive and motor impairments contribute to reduced quality of life and are more common. Regional cerebellar volume is associated with performance across these domains and sex hormones may influence this volume. Examining sex differences in regional cerebellar volume in conjunction with age, and in the context of reproductive stage stands to improve our understanding of cerebellar aging and pathology.

**Methods:** Data from 530 healthy adults (ages 18-88; 49% female) from the Cambridge Centre for Ageing and Neuroscience database were used here. CERES was utilized to assess lobular volume in T1-weighted images. We examined sex differences in adjusted regional cerebellar volume while controlling for age. A subgroup (n = 354, 50% female) was used to assess group differences in female reproductive stages as compared to age-matched males.

**Results:** Sex differences in adjusted volume were seen across most anterior and posterior cerebellar lobules. The majority of cerebellar regions had significant linear relationships with age in males and females. However, there were no interactions between sex and reproductive stage groups (i.e., female reproductive stage did not display a relationship with regional cerebellar volume).

**Discussion:** We found sex differences in volume across much of the cerebellum, linear associations with age, and did not find an effect of female reproductive stage on regional cerebellar volume. Longitudinal investigation into hormonal influences on cerebellar structure and function is warranted as hormonal changes with menopause may impact structure over time.

## Introduction

Aging is associated with deficits in cognitive and motor functioning (Harada, Love, & Triebel, 2013; Lezak et al., 2004; Siedler et al., 2010). These deficits can interfere with activities of daily living and general independence in advanced age (Kluger et al., 1997; Schaefer & Schumacher, 2011). Furthermore, motor deficits such as gait speed have been linked to cognitive decline (e.g., memory loss), and in some cases can predict cognitive deficits (Kikkert et al., 2016; Marquis et al., 2002; Savica et al., 2017). While the cerebellum has long established associations with motor function, there is also a large and growing literature implicating it in cognition (Balsters, Whelan, Robertson, & Ramnani, 2013; Bernard et al., 2020; Chen & Desmond, 2005; Stoodley & Schmahmann, 2009). Topographical organization based on functional imaging has implicated Crus I, Crus II, and Lobules VI-VIIB in cognitive processes, and Lobules I-V, Lobules VIIIA-VIIIB in motor processes (Guell & Schmahmann, 2020; King et al., 2019; Stoodley & Schmahmann, 2009; Stoodley, Valera, & Schmahmann, 2012). Notably, recent studies suggest these relationships are not strictly confined to lobular boundaries and more closely resemble a continuum (Guell & Schmahmann, 2020; King et al., 2019). However, older adult cerebella show functionally relevant volumetric differences (Bernard & Seidler, 2013; Miller et al., 2013). Additionally, differences in functional connectivity between the cerebellum and cortex during resting state are associated with cognitive decline in older adults (Bernard & Siedler, 2013a; de Dieu Uwisengeyimana, et al., 2020). Thus, advanced age impacts the cerebellum, which contributes to cognitive and motor processes.

The extant literature has demonstrated global cerebellar neurodegeneration in advanced age (Jernigan et al. 2001; Hulst et al. 2015; Raz et al. 1998, 2010). Yet the regional volumetric trajectory of the cerebellum across the adult lifespan has been less studied. Given the functional topography of the cerebellum, characterization of regional cerebellar structure in normal aging may help contextualize functional changes. To date, several studies have shown age differences in regional cerebellar volume (Bernard & Seidler, 2013; Koppelmans et al., 2017; Han et al., 2020). Cross-sectional analyses of these volumes in relation to age demonstrate age differences in certain cerebellar regions across studies (i.e., right Lobules I-III, bilateral Lobules V and VI, bilateral Crus I, and left Crus II) (Bernard & Seidler, 2013; Han et al., 2020; Koppelmans et al., 2017). However, in each study the age range of the sample varied (Bernard & Seidler, 2013; Han et al., 2020; Koppelmans et al., 2017), and to this point, no single analysis has looked at the totality of adulthood (from age 18 up) in one sample. This work stands to enhance our understanding of the aging process and emergent pathology, while also providing a more complete picture of relationships between cerebellar volume and age across the adult lifespan.

Sex is also an important factor to consider with respect to cerebellar aging. Studies evaluating global measures of cerebellar volume across adulthood demonstrated greater white matter volume in the posterior cerebellum of males, greater hemispheric volume in males, and greater volumetric atrophy in aging for males (Geng et al., 2016; Raz et al., 2001; Xu et al., 2000). However, a recent meta-analysis reported mixed results with respect to sex differences in cerebellar volume (Eliot et al., 2021). In a study of young adults, females had larger volumes in bilateral Crus II and Vermis VI while males exhibited larger right lobule V, bilateral lobules VIIIA/B, and Vermis VIIIA/B (Steele & Chakravaty, 2018). A cross-sectional evaluation of older adults found significant sex differences in Vermis IX and X, right Lobules I-III, and bilateral Lobules VIIIA, while longitudinal analyses of the same sample demonstrated significant sex differences over time in left Lobule VI and right Crus II (Han et al., 2020). Luft and colleagues boasted a wider range of adults (ages 19-73) in which females had greater adjusted volume in Lobules VI-VII and the Superior Posterior Lobe than males (Luft et al., 1999). Thus far, studies examining sex differences in cerebellar volume have demonstrated considerable inconsistencies and varying age groups/ranges, warranting more nuanced investigation.

Prior inconsistencies in sex differences may be due, at least in part, to hormonal factors which play a role in modulating cerebellar structure and function. For instance, a typical female menstrual cycle is characterized by a 12-fold increase in progesterone and 80-fold increase in estrogen (Taylor, Pritschet, Yu, & Jacobs, 2019). Females experience significant physiological shifts due to the menopausal transition (i.e., 90% decrease in ovarian estradiol production) and decreased estrogen has been associated with increased risk of AD pathogenesis in the aging brain (Buckler, 2005; Harlow et al. 2012; Hedges et al., 2012; Morrison & Baxter, 2012; Vest & Pike, 2013). Furthermore, endocrine aging (as it relates to the menopausal transition) has greater effects than chronological aging on the female cortex (Mosconi et al., 2017) which raises the question whether there are hormonal influences on regional cerebellar structure in healthy aging. Sex hormones have demonstrated influences on brain structure and function (e.g., cortical connectivity, subcortical connectivity, and within-network coherence) especially in regions dense with estrogen receptors in the context of both endogenous hormone levels and exogenous sex hormone manipulations (Peper et al., 2011; Pritschet et al., 2020; Taylor, Pritschet, Yu, & Jacobs, 2019). While there are estrogen receptors throughout the brain, the cerebellum is one of the few regions dense with these receptors (Boyle et al., 2021). However, the influence of sex hormones specifically on the cerebellum are less studied and warrant further investigation (Hedges et al., 2012; Peper et al., 2011). Given the scope of hormonal influences and the considerable variance of sex hormone levels in reproductive aging, different reproductive stages in adult females may be critical periods of change in cerebellar structure and exhibit different patterns of cerebellar volume across the female adult lifespan.

Here, we examined lobular cerebellar volume across the adult lifespan (in cross-sectional data) to investigate associations with age, sex differences, and female reproductive stage. We tested the following hypotheses: 1) there would be sex differences in volumes of cerebellar lobules when controlling for age; 2) age-related patterns of regional cerebellar volume would diverge between sexes across the adult lifespan; and 3) volumetric differences would be present when looking at females grouped by reproductive stage. Establishing whether there are sex differences in regional cerebellar volume more generally and at different female reproductive stages will contribute to our understanding of the aging cerebellum and in turn may serve as a foundation for work investigating behavioral decline, including that with clinical significance (e.g., Alzheimer’s Disease).

## Methods

### Participants

Data used in these analyses are from the Cambridge Centre for Ageing and Neuroscience (CamCAN) repository which collected epidemiological, cognitive, genetic, and neuroimaging data to examine normative neurocognitive development (available at http://www.mrc-cbu.cam.ac.uk/datasets/camcan/) (Shafto et al. 2014; Taylor et al. 2017). The Cam-CAN sample was population-based and originally consisted of 708 healthy adults aged 18-88. Magnetic resonance imaging (MRI) scans of the brain were acquired at the Medical Research Council Cognition & Brain Sciences Unit in Cambridge, UK. Here, we used the high-resolution T1 anatomical scans collected on a 3T Siemens TIM Trio System. Participants with T1 images displaying poor resolution, notable artifacts, or error messages related to image quality during segmentation processing were excluded from this study (n = 178). For a more detailed account of this study please refer to Shafto and colleagues’ description (2014) and Taylor and colleagues (2017).

This study developed two subsets of the Cam-CAN sample for analyses due to analytic redundancies between reproductive stage and age. The first subset of participants (n = 530, 49% female) were utilized to evaluate associations between age and regional cerebellar volume in males and females across adult lifespan, while the second subset of participants (n = 354, 50% female) examined females against age-matched males in each reproductive stage (further description of both subsets below).

### Reproductive Stage Categorization

As part of the Cam-CAN data repository, female participants reported the number of days since their last menstrual period, length of their menstrual cycle in days, and the age when menstrual periods ceased. This self-report data was used to categorize these females into reproductive groups based on STRAW+10 guidelines (Harlow et al., 2012). Females were categorized into the following stages: reproductive, menopausal transition (i.e., perimenopause), and postmenopause with further temporal specifications of early and late subgroups (Harlow et al., 2012). The reproductive stage is characterized by a regular menstrual cycle which may include subtle changes in flow length; participant indication of a menstrual cycle 35 days or less were placed in this group (Harlow et al., 2012). In the absence of self-report data, females were categorized *a priori* into reproductive groups by age as follows: reproductive ≤ 36 years, perimenopause 40-50 years, early postmenopause 55-70, late postmenopause ≥ 71 years. Five females lacking self-report data from ages 50-55 were excluded due to the high variability of reproductive stage during that time period. Perimenopause is categorized via an inconsistent cycle with persistent ≥ 7 day difference in length of consecutive cycles or intervals of amenorrhea of greater than 60 days. Individuals that self-reported a menstrual cycle greater than 35 days or greater than 60 days since their last menstrual period, but not yet in menopause were allocated to this group (Harlow et al., 2012). The early postmenopausal stage describes the 2-6 year period after the menstrual cycle ceases; females whose age was within 2-6 years of the reported age when menstrual periods stopped were included in this group (Harlow et al., 2012). Lastly, late postmenopause is characterized by the remaining female lifespan; participants with an age greater than 6 years from their reported age of last menstrual cycle were assigned to this category (Harlow et al., 2012).

After categorization of reproductive stages, matching females to same aged males in each reproductive stage was employed to control for effects of age and to differentiate effects of sex. As reproductive stage and age are interdependent, age as a variable created redundancies in the analyses which necessitated the use of age-matched males. Additionally, early visualizations of these data indicated outliers in multiple regions. As our analyses compare group averages, outliers could skew our results. Our process required exclusion criteria which was completed in four steps. In the first step, outliers were removed. Outliers were categorized as any value three standard deviations above or below the mean within each reproductive stage group. The mean and standard deviation for each region of each group was calculated in excel and a logical ‘if’ formula identified outliers. Each outlier was manually removed along with an age-matched participant. Next, female participants were manually matched via exact age in years to a male. The following steps were conducted to further exclude participants when the number of females and males of matching age were unequal. The number of females in each reproductive stage group notably varied between groups (i.e., final group breakdown: reproductive (n = 126), perimenopause (n = 54), early postmenopause (n = 56), and late postmenopause (n = 118)). Thus, the third step prioritized categorizing an age-matched male into the smallest reproductive stage groups first in an effort to more robustly represent each reproductive stage. In cases where there were more females than males or vice versa, a fourth step eliminated the males or females with the highest signal-to-noise ratio (SNR) on their T1 scans. For example, if there was one female at age 40 and three males at age 40, the male with the lowest SNR would remain in the dataset and be categorized into the same reproductive stage as that female; whereas the two 40-year old males with higher SNR would be excluded. Thus, this step included only participants with the best SNR, but this was only used if the previous two steps left an unequal number of females and males. This exclusion process reduced the sample size of these analyses from n = 530 to n = 354.

### Neuroimaging Data

These data were collected on a 3T Siemens TIM Trio System. Imaging parameters for T1 scans of the brain were as follows: 3D MPRAGE, TR=2250ms, TE=2.99ms, TI=900ms; FA=9 deg; FOV=256×240×192mm; 1mm isotropic; GRAPPA=2; TA=4mins 32s.

Lobular cerebellar volume was computed using CERES (Romero et al., 2017). CERES is an automated cerebellum segmentation method which only requires a standard resolution T1- weighted magnetic resonance image (Romero et al., 2017). Of note, CERES lobular segmentation divides the vermis down its midline into left and right regions, including vermis volumes within each lobule. As such, vermal subregions were not considered separately in this analysis. CERES exhibited the highest accuracy in the majority of regions examined in a comparison of three automated parcellation programs and had the greatest intra-rater accuracy between automated segmentation and expert manual segmentation (Romero et al., 2017). When compared to other cerebellar segmentation algorithms, CERES demonstrated the best performance across the majority of regions examined (Carass et al., 2018). To use CERES, T1- weighted nifti files were uploaded directly to the volBrain website (https://www.volbrain.upv.es/index.php). The CERES algorithm executes preprocessing on the files (i.e., denoising, bias field correction using the N4 algorithm, MNI affine registration, cropping the cerebellum, non-linear registration estimation using Advanced Normalizations Tools (ANTs), and local intensity normalization) and labeling cerebellar voxels (via three steps: 1) non-local patch-based local fusion, 2) the PatchMatch algorithm, and 3) Optimized PatchMatch) (Barnes, Shechtman, Finkelstein, & Goldman, 2009; Romero et al., 2017; Ta, Giraud, Collins, & Coupé, 2014).

### Statistical Analyses

The variables, statistical packages, and manual corrections were consistent across all analyses. All analyses used adjusted volumes that were corrected for estimated intracranial volume, as volume is also proportional to overall head and body size (Buckner et al., 2004). This was calculated as the ratio of lobular volume to estimated intracranial volume (region in cm^3^/estimated intracranial volume) via CERES’ software and included in the output. Analyses of covariance (ANCOVAs) were completed using the anova function from the default ‘stats’ package in R (v4.0.5, R Core Team, 2021) which determined beta coefficients, degrees of freedom, F-values, and p-values; sjstats and pwr packages were used for **η**^2^ and partial **η**^2^ values (v0.18.1, Lüdecke, 2018). A multivariate analysis of variance (MANOVA) was completed using the manova function from the default ‘stats’ package in R (v4.0.5, R Core Team, 2021) which determined Pillai’s trace, degrees of freedom, p-values, and approximated F-values. Linear and quadratic regressions were performed using the lm function from the default ‘stats’ package in R (v4.0.5, R Core Team, 2021) which determined beta coefficients, degrees of freedom, F-values, p-values, R^2^, and adjusted R^2^. In each analysis Bonferroni correction was manually applied to account for multiple comparisons (.05/24 for significance at *p* = 0.002083).

### Regional Cerebellar Sex Difference Analyses

ANCOVAs were conducted to investigate sex differences in right and left cerebellar lobules while controlling for age. Violin plots were created using the ggplot2 package (Figure 1; v3.3.3; Wickham, 2016).

**Figure 1.**
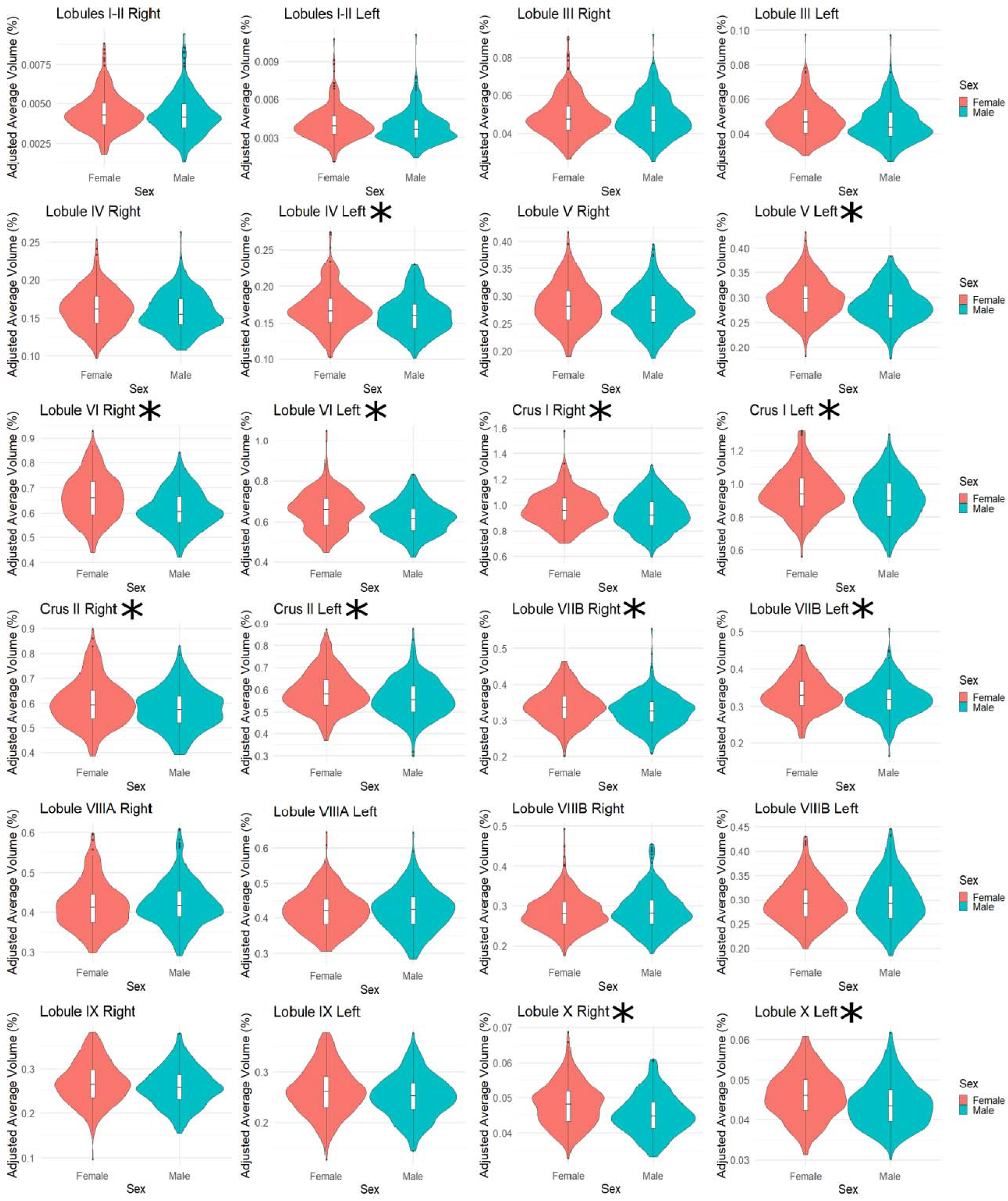
Sex Difference Adjusted Volume Distributions in Cerebellar Regions. Violin plots demonstrating corected volume for each lobule in males and females across all participants. The curved edges depict the distribution of volume in these participants and box plots indicate the mean and interquartile ranges. The asterisk depicts areas that demonstrated a statistically significant difference by sex when controlling for age after Bonferroni correction (*p*<.002083).

### Linear and Quadratic Age-Volume Associations by Sex

Age-volume associations were evaluated via linear regression with age as the predictor and right and left adjusted regional cerebellar volumes as the outcome in males and females separately. These analyses were also completed using a quadratic age variable (Age + I(Age^2)) as the predictor to explore whether adjusted regional volume is fit with a quadratic function rather than linear across the lifespan, given prior work suggesting non-linear relationships between volume and age (Bernard et al., 2015). Akaike’s An Information Criterion (AIC) (Sakamoto, Ishiguro, & Kltagawa, 1986) compared fit between linear and quadratic models with a requirement that model value must differ by 10 to be considered a superior fit (Bohon & Welch, 2021; Burnham & Anderson, 2004). AIC was calculated by the default ‘stats’ package in R (v4.0.5, R Core Team, 2021). The stargazer package (v5.2.2; Hlavac, 2018) was utilized to create Tables II-V. The ggplot2 package was used to create linear plots (Figure 2; v3.3.3; Wickham, 2016).

**Figure 2.**
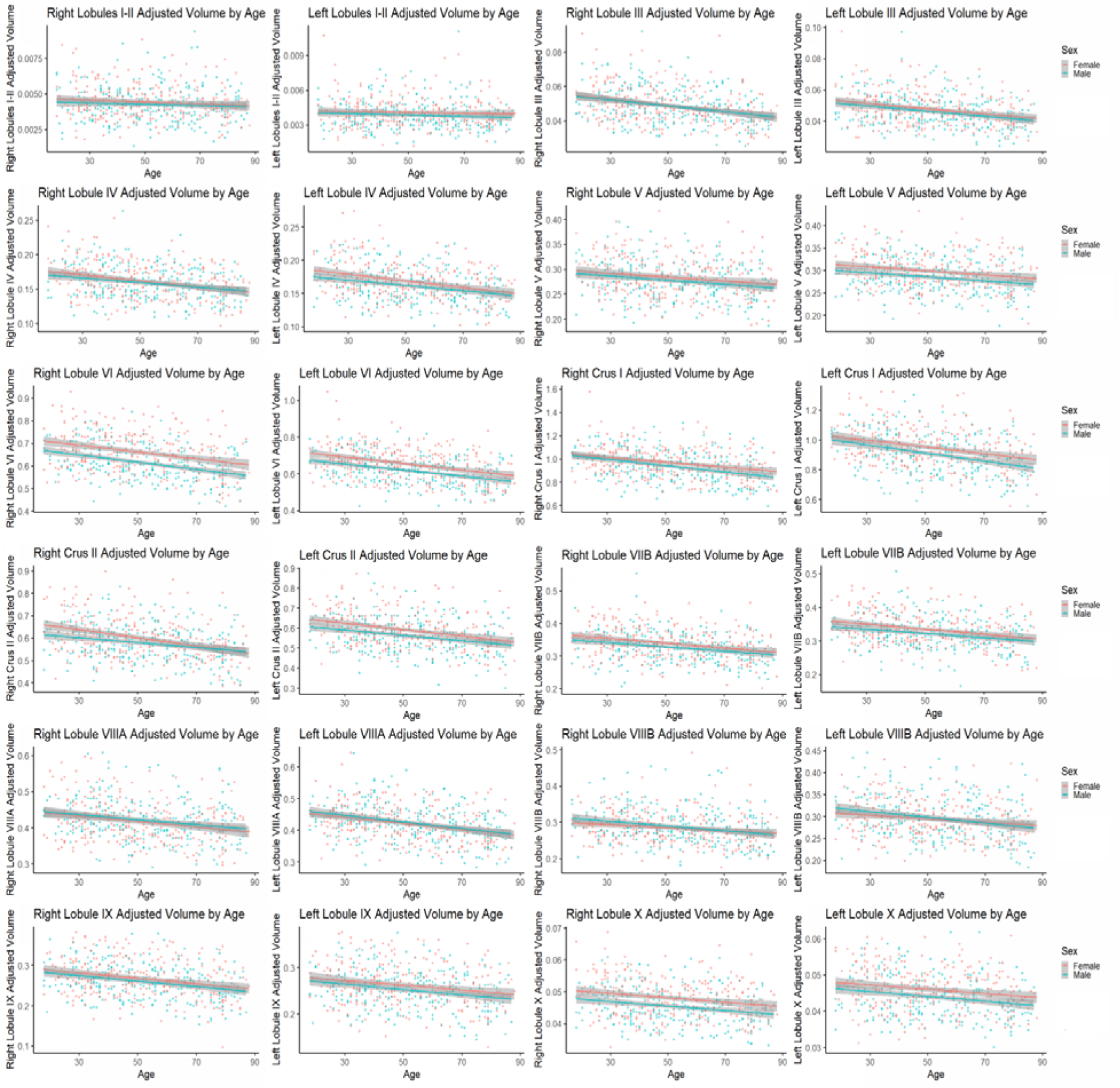
Linear Age Associations with Adjusted Volume in Cerebellar Regions by Sex. Linear age-volume relationships by sex in each cerebellar region examined. The gray superimposed on each colored line depicts the 95% confidence interval for adjusted volume in each sex.

### Reproductive Status Analyses

To evaluate differences in adjusted regional cerebellar volume by reproductive stage, we used a MANOVA for right and left cerebellar regions in reproductive (n = 146), perimenopause (n = 56), early postmenopause (n = 58), and late postmenopause (n = 134) stages in females as compared to age-matched males (Harlow et al., 2012). Reproductive stage (4 groups) and sex (2 groups) were categorical predictors and adjusted regional cerebellar volume was the outcome in this 4 × 2 MANOVA. Figures depicting group descriptives were created using default jamovi descriptive plots (Version 2.0.0.0; The jamovi project, 2021; Figure 3).

**Figure 3.**
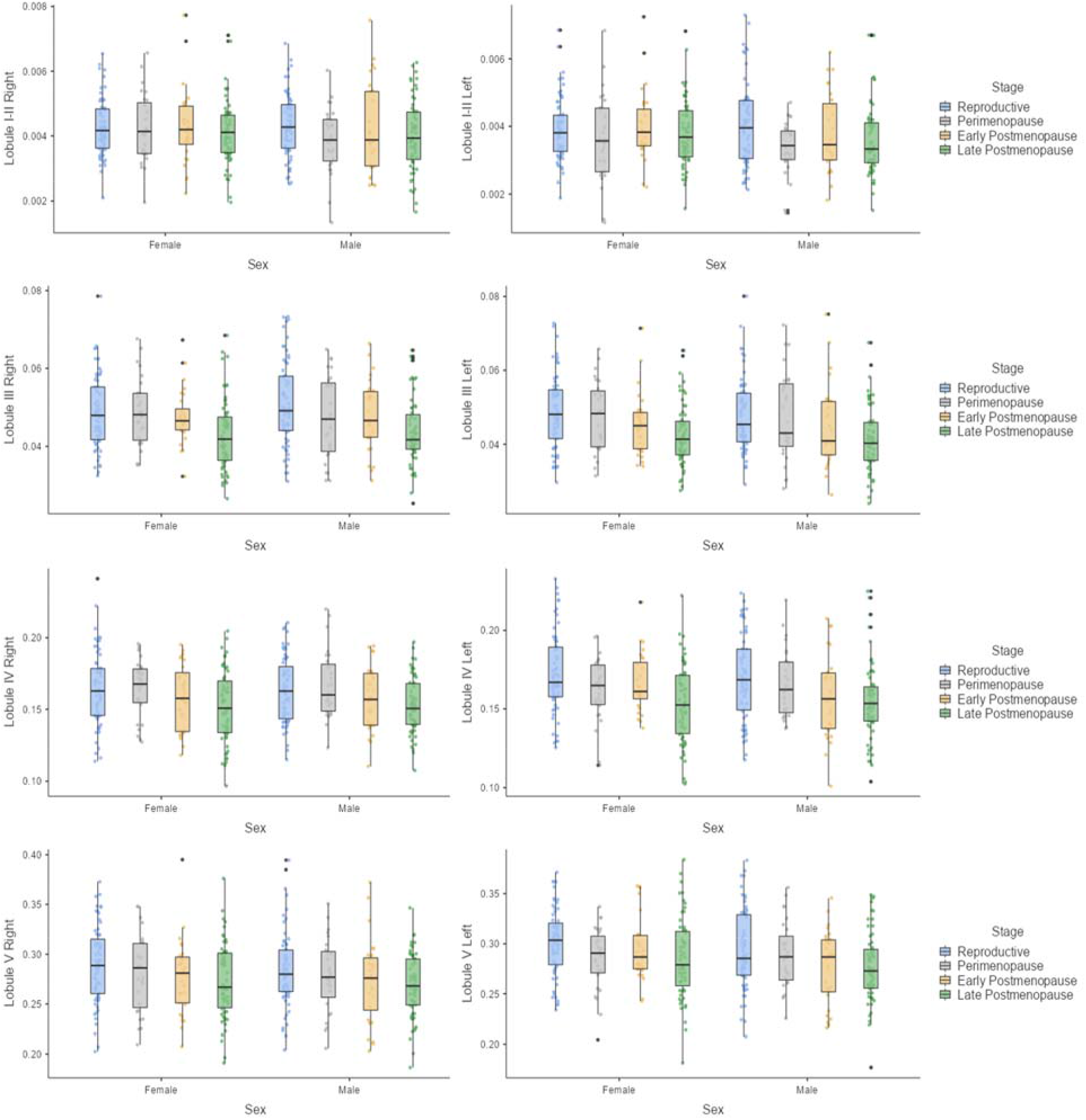
Reproductive Stage Associations with Adjusted Volume in Cerebellar Regions by Sex. Mean, interquartile range, and adjusted volume distribution for age-matched males and females in categorized female reproductive stages. There were no significant interactions.

## Results

The sum of squares, mean squared error, F-statistic, p-value, η^2^, and partial η^2^ are presented for each cerebellar region in Table I. The estimated **β** coefficients, standard error, R^2^, adjusted R^2^, residual standard error, and F-statistic are shown in Tables II-V with 3 asterisks indicating a significant *p*-value after Bonferroni correction.

**Table I.** Sex Differences in Adjusted Cerebellar Volume.

| **Sex Differences in Adjusted Cerebellar Volume** |  |  |  |  |  |  |
| --- | --- | --- | --- | --- | --- | --- |
| **Cerebellar Region** | **Sum of Squares** | **Mean Square** | **F Statistic (df = 1; 527)** | **p-value** | **$\eta^2$** | **Partial $\eta^2$** |
| **I-II right** | 0.00000 | 0.00000 | 2.881 | 0.090 | 0.005 | 0.005 |
| **I-II left** | 0.00001 | 0.00001 | 5.615 | 0.018* | 0.01 | 0.011 |
| **III right** | 0.00005 | 0.00005 | 0.553 | 0.458 | 0.00095 | 0.001 |
| **III left** | 0.00055 | 0.00055 | 5.408 | 0.020* | 0.009 | 0.01 |
| **IV right** | 0.00200 | 0.00200 | 2.757 | 0.097 | 0.005 | 0.005 |
| **IV left** | 0.00800 | 0.00800 | 13.051 | < .001*** | 0.022 | 0.024 |
| **V right** | 0.00800 | 0.00800 | 5.759 | 0.017* | 0.01 | 0.011 |
| **V left** | 0.02700 | 0.02700 | 19.773 | < .001*** | 0.035 | 0.036 |
| **VI right** | 0.31600 | 0.31600 | 50.344 | < .001*** | 0.078 | 0.087 |
| **VI left** | 0.21100 | 0.21100 | 33.215 | < .001*** | 0.052 | 0.059 |
| **Crus I right** | 0.17000 | 0.17000 | 11.046 | < .001*** | 0.018 | 0.021 |
| **Crus I left** | 0.30400 | 0.30400 | 19.027 | < .001*** | 0.031 | 0.035 |
| **Crus II right** | 0.07600 | 0.07600 | 10.999 | < .001*** | 0.019 | 0.02 |
| **Crus II left** | 0.13400 | 0.13400 | 19.457 | < .001*** | 0.032 | 0.036 |
| **VIIB right** | 0.02300 | 0.02300 | 11.621 | < .001*** | 0.02 | 0.022 |
| **VIIB left** | 0.02700 | 0.02700 | 12.899 | < .001*** | 0.022 | 0.024 |
| **VIIIA right** | 0.00200 | 0.00200 | 0.567 | 0.452 | 0.001 | 0.001 |
| **VIIIA left** | 0.00040 | 0.00040 | 0.131 | 0.718 | 0.00022 | 0.00025 |
| **VIIIB right** | 0.00200 | 0.00200 | 0.791 | 0.374 | 0.001 | 0.002 |
| **VIIIB left** | 0.00057 | 0.00057 | 0.275 | 0.601 | 0.0005 | 0.00052 |
| **IX right** | 0.01100 | 0.01100 | 6.015 | 0.015* | 0.01 | 0.011 |
| **IX left** | 0.01600 | 0.01600 | 8.553 | 0.004** | 0.015 | 0.016 |
| **X right** | 0.00100 | 0.00100 | 30.501 | < .001*** | 0.052 | 0.055 |
| **X left** | 0.00060 | 0.00060 | 18.701 | < .001*** | 0.033 | 0.034 |

*Note. Significance levels. \* = $p \leq 0.05$ , \*\* = $p \leq 0.01$ , \*\*\* = $p \leq 0.001$*

### Regional Cerebellar Sex Differences

ANCOVAs revealed significant sex differences in adjusted volume for the following regions: left Lobule IV, left Lobule V, left and right Lobule VI, left and right Lobule VIIB, left and right Lobule X, left and right Crus I, and left and right Crus II, when controlling for age and following Bonferroni correction [F(1,527) > 10.999, *p*<.001, for detailed results, please see Table I]. There were no significant sex differences in any other regions (*p*>.002083). Figure 1 displays the results of these analyses.

### Linear and Quadratic Age-Volume Associations by Sex

Males and females were analyzed separately to ascertain whether adjusted regional cerebellar volume is associated with age over the adult lifespan. Linear regressions in males with age as the predictor and adjusted volume as the outcome revealed a significant relationship with age in the following areas: bilateral Lobules III, IV, V, VI, VIIB, VIIIA, VIIIB, IX, X, and bilateral Crus I and Crus II [F(1,266) > 11.960, *p<.001*, detailed results are presented in Table II and Figure 2], after Bonferroni correction. Bilateral Lobules I-II were the only regions that did not demonstrate a significant linear relationship with age in either sex (*p*>.05).

**Table II.**
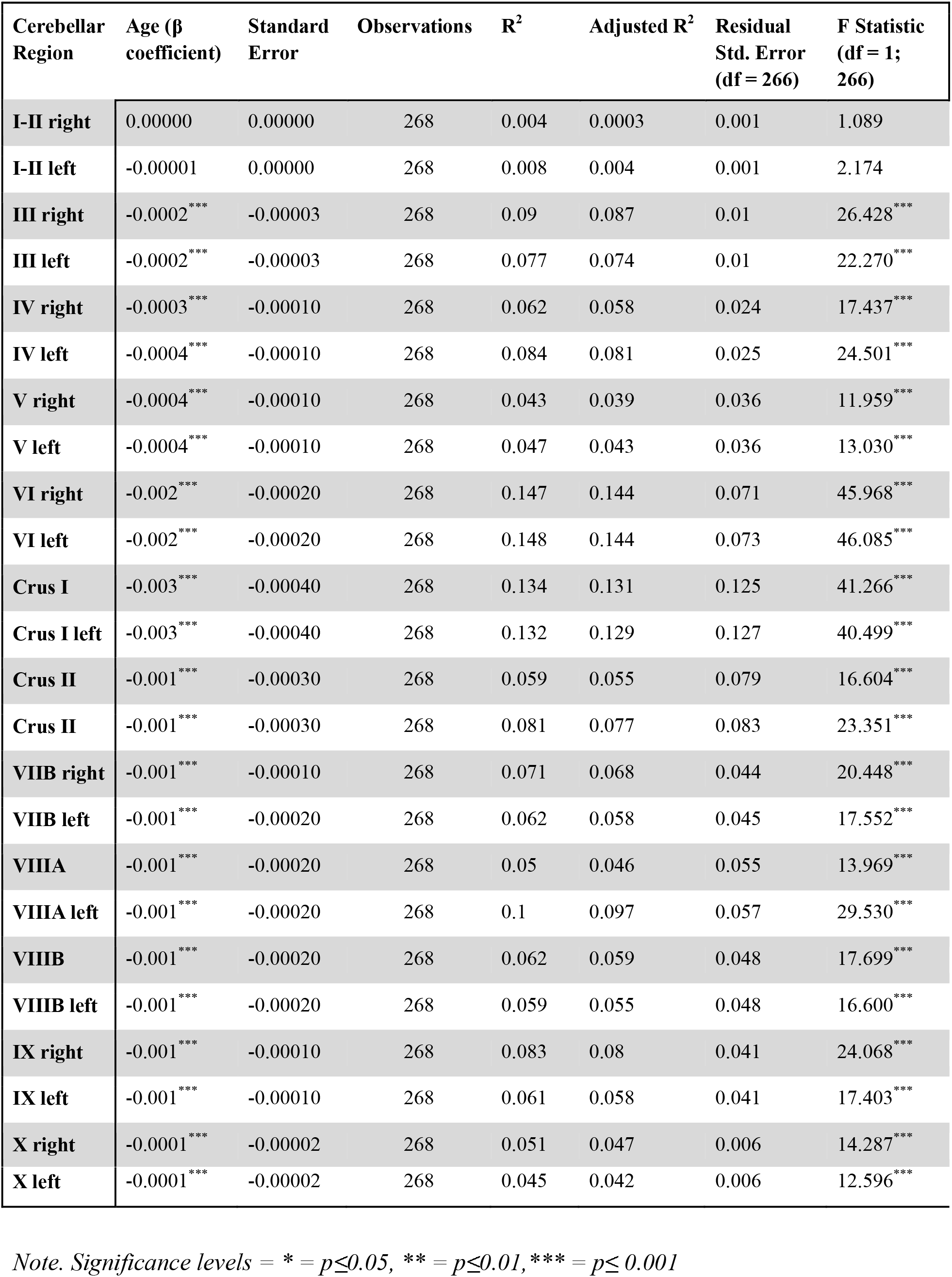
Linear Relationship with Age and Adjusted Cerebellar Volume in Males.

| Linear Relationship with Age and Adjusted Cerebellar Volume in Males |  |  |  |  |  |  |  |
| --- | --- | --- | --- | --- | --- | --- | --- |
| Cerebellar Region | Age ($\beta$ coefficient) | Standard Error | Observations | $R^2$ | Adjusted $R^2$ | Residual Std. Error (df = 266) | F Statistic (df = 1; 266) |
| I-II right | 0.00000 | 0.00000 | 268 | 0.004 | 0.0003 | 0.001 | 1.089 |
| I-II left | -0.00001 | 0.00000 | 268 | 0.008 | 0.004 | 0.001 | 2.174 |
| III right | -0.0002*** | -0.00003 | 268 | 0.09 | 0.087 | 0.01 | 26.428*** |
| III left | -0.0002*** | -0.00003 | 268 | 0.077 | 0.074 | 0.01 | 22.270*** |
| IV right | -0.0003*** | -0.00010 | 268 | 0.062 | 0.058 | 0.024 | 17.437*** |
| IV left | -0.0004*** | -0.00010 | 268 | 0.084 | 0.081 | 0.025 | 24.501*** |
| V right | -0.0004*** | -0.00010 | 268 | 0.043 | 0.039 | 0.036 | 11.959*** |
| V left | -0.0004*** | -0.00010 | 268 | 0.047 | 0.043 | 0.036 | 13.030*** |
| VI right | -0.002*** | -0.00020 | 268 | 0.147 | 0.144 | 0.071 | 45.968*** |
| VI left | -0.002*** | -0.00020 | 268 | 0.148 | 0.144 | 0.073 | 46.085*** |
| Crus I | -0.003*** | -0.00040 | 268 | 0.134 | 0.131 | 0.125 | 41.266*** |
| Crus I left | -0.003*** | -0.00040 | 268 | 0.132 | 0.129 | 0.127 | 40.499*** |
| Crus II | -0.001*** | -0.00030 | 268 | 0.059 | 0.055 | 0.079 | 16.604*** |
| Crus II | -0.001*** | -0.00030 | 268 | 0.081 | 0.077 | 0.083 | 23.351*** |
| VIIB right | -0.001*** | -0.00010 | 268 | 0.071 | 0.068 | 0.044 | 20.448*** |
| VIIB left | -0.001*** | -0.00020 | 268 | 0.062 | 0.058 | 0.045 | 17.552*** |
| VIIIA | -0.001*** | -0.00020 | 268 | 0.05 | 0.046 | 0.055 | 13.969*** |
| VIIIA left | -0.001*** | -0.00020 | 268 | 0.1 | 0.097 | 0.057 | 29.530*** |
| VIIIB | -0.001*** | -0.00020 | 268 | 0.062 | 0.059 | 0.048 | 17.699*** |
| VIIIB left | -0.001*** | -0.00020 | 268 | 0.059 | 0.055 | 0.048 | 16.600*** |
| IX right | -0.001*** | -0.00010 | 268 | 0.083 | 0.08 | 0.041 | 24.068*** |
| IX left | -0.001*** | -0.00010 | 268 | 0.061 | 0.058 | 0.041 | 17.403*** |
| X right | -0.0001*** | -0.00002 | 268 | 0.051 | 0.047 | 0.006 | 14.287*** |
| **X left** | -0.0001<sup>***</sup> | -0.00002 | 268 | 0.045 | 0.042 | 0.006 | 12.596<sup>***</sup> |

*Note. Significance levels = \* = $p \leq 0.05$ , \*\* = $p \leq 0.01$ , \*\*\* = $p \leq 0.001$*

In females, there was a significant linear relationship between age and adjusted regional cerebellar volume in the following areas: bilateral Lobules III, IV, VI, VIIB, VIIIA, IX, left Lobule V, right Lobule X, and bilateral Crus I and Crus II [F(1,260) > 12.080, *p*<.001] after Bonferroni correction. Detailed results are presented in Table III and Figure 2.

**Table III.** Linear Relationship with Age and Adjusted Cerebellar Volume in Females.

|  |  |  |  |  |  |  |  |
| --- | --- | --- | --- | --- | --- | --- | --- |
| Cerebellar Region | Age ($\beta$ coefficient) | Standard Error | Observations | R<sup>2</sup> | Adjusted R<sup>2</sup> | Residual Std. Error (df = 260) | F Statistic (df = 1; 260) |
| I-II right | -0.00001* | 0.00000 | 262 | 0.011 | 0.007 | 0.001 | 2.881* |
| I-II left | 0.00000 | 0.00000 | 262 | 0.002 | -0.002 | 0.001 | 0.602 |
| III right | -0.0002*** | -0.00003 | 262 | 0.104 | 0.101 | 0.01 | 30.215*** |
| III left | -0.0002*** | -0.00003 | 262 | 0.089 | 0.085 | 0.01 | 25.314*** |
| IV right | -0.0004*** | -0.0001 | 262 | 0.098 | 0.095 | 0.025 | 28.274*** |
| IV left | -0.0005*** | -0.0001 | 262 | 0.107 | 0.103 | 0.026 | 31.060*** |
| V right | -0.0004*** | -0.0001 | 262 | 0.032 | 0.028 | 0.039 | 8.458** |
| V left | -0.0004*** | -0.0001 | 262 | 0.044 | 0.041 | 0.038 | 12.076*** |
| VI right | -0.001*** | -0.0003 | 262 | 0.093 | 0.089 | 0.087 | 26.544*** |
| VI left | -0.002*** | -0.0003 | 262 | 0.128 | 0.125 | 0.086 | 38.213*** |
| Crus I | -0.002*** | -0.0004 | 262 | 0.091 | 0.087 | 0.123 | 25.916*** |
| Crus I left | -0.002*** | -0.0004 | 262 | 0.101 | 0.097 | 0.126 | 29.175*** |
| Crus II | -0.002*** | -0.0003 | 262 | 0.129 | 0.125 | 0.087 | 38.357*** |
| Crus II left | -0.002*** | -0.0003 | 262 | 0.111 | 0.108 | 0.084 | 32.573*** |
| VIIB right | -0.001*** | -0.0002 | 262 | 0.084 | 0.08 | 0.046 | 23.699*** |
| VIIB left | -0.001*** | -0.0002 | 262 | 0.081 | 0.078 | 0.046 | 23.069*** |
| VIIIA | -0.001*** | -0.0002 | 262 | 0.058 | 0.054 | 0.057 | 15.950*** |
| VIIIA left | -0.001*** | -0.0002 | 262 | 0.102 | 0.099 | 0.053 | 29.635*** |
| VIIIB | -0.0004*** | -0.0001 | 262 | 0.029 | 0.025 | 0.043 | 7.795** |
| VIIIB left | -0.0004*** | -0.0001 | 262 | 0.028 | 0.024 | 0.042 | 7.369** |
| **IX right** | -0.001*** | -0.0002 | 262 | 0.07 | 0.066 | 0.046 | 19.435*** |
| **IX left** | -0.001*** | -0.0002 | 262 | 0.045 | 0.041 | 0.045 | 12.280*** |
| **X right** | -0.0001*** | -0.00002 | 262 | 0.045 | 0.041 | 0.006 | 12.280*** |
| **X left** | -0.0001*** | -0.00002 | 262 | 0.035 | 0.031 | 0.006 | 9.332** |

*Note. Significance levels = \* = $p \leq 0.05$ , \*\* = $p \leq 0.01$ , \*\*\* = $p \leq 0.001$*

Bilateral Lobules I-II, right Lobule V, bilateral Lobules VIIIB, and left Lobule X did not demonstrate a significant linear relationship with age in females (*p>.002083*). The quadratic variables in regressions were not significant for any regions in males or females (*p*>.05 and *p*>.002, respectively; please see Tables IV and V).

**Table IV.** Quadratic Relationship with Age and Adjusted Cerebellar Volume in Males.

|  |  |  |  |  |  |  |  |
| --- | --- | --- | --- | --- | --- | --- | --- |
| **Cerebellar Region** | **Quadratic Age ($\beta$ coefficient)** | **Standard Error** | **Observations** | **R<sup>2</sup>** | **Adjusted R<sup>2</sup>** | **Residual Std. Error (df = 265)** | **F Statistic (df = 2; 265)** |
| **I-II right** | 0.00000 | 0.00000 | 268 | 0.005 | -0.002 | 0.001 | 0.677 |
| **I-II left** | 0.00000 | 0.00000 | 268 | 0.014 | 0.007 | 0.001 | 1.931 |
| **III right** | 0.00000 | 0.00000 | 268 | 0.097 | 0.09 | 0.01 | 14.163*** |
| **III left** | -0.00000* | 0.00000 | 268 | 0.089 | 0.083 | 0.01 | 13.012*** |
| **IV right** | 0.00000 | 0.00000 | 268 | 0.062 | 0.055 | 0.024 | 8.784*** |
| **IV left** | 0.00000 | 0.00000 | 268 | 0.086 | 0.079 | 0.025 | 12.443*** |
| **V right** | 0.00000 | -0.00001 | 268 | 0.043 | 0.036 | 0.036 | 6.022*** |
| **V left** | -0.00001 | -0.00001 | 268 | 0.051 | 0.044 | 0.036 | 7.144*** |
| **VI right** | 0.00000 | -0.00001 | 268 | 0.148 | 0.141 | 0.071 | 22.940*** |
| **VI left** | 0.00001 | -0.00001 | 268 | 0.151 | 0.145 | 0.073 | 23.591*** |
| **Crus I right** | -0.00001 | -0.00002 | 268 | 0.135 | 0.128 | 0.125 | 20.668*** |
| **Crus I left** | 0.00001 | -0.00002 | 268 | 0.133 | 0.126 | 0.127 | 20.254*** |
| **Crus II right** | -0.00002 | -0.00002 | 268 | 0.065 | 0.058 | 0.079 | 9.169*** |
| **Crus II left** | -0.00001 | -0.00002 | 268 | 0.081 | 0.074 | 0.083 | 11.687*** |
| **VIIB right** | -0.00001 | -0.00001 | 268 | 0.079 | 0.072 | 0.044 | 11.378*** |
| **VIIB left** | -0.00001 | -0.00001 | 268 | 0.064 | 0.057 | 0.045 | 9.002*** |
| **VIIIA right** | 0.00000 | -0.00001 | 268 | 0.05 | 0.043 | 0.055 | 7.036*** |
| **VIIIA left** | -0.00001 | -0.00001 | 268 | 0.105 | 0.098 | 0.057 | 15.559*** |
| **VIIIB right** | -0.00001 | -0.00001 | 268 | 0.064 | 0.057 | 0.048 | 9.067*** |
| **VIIIB left** | -0.00001 | -0.00001 | 268 | 0.06 | 0.053 | 0.048 | 8.512*** |
| **IX right** | -0.00001 | -0.00001 | 268 | 0.088 | 0.081 | 0.041 | 12.775*** |
| **IX left** | -0.00001 | -0.00001 | 268 | 0.067 | 0.06 | 0.041 | 9.548*** |
| **X right** | 0.00000 | 0.00000 | 268 | 0.051 | 0.044 | 0.006 | 7.157*** |
| **X left** | 0.00000 | 0.00000 | 268 | 0.049 | 0.042 | 0.006 | 6.887*** |

*Note. Significance levels = \* = $p \leq 0.05$ , \*\* = $p \leq 0.01$ , \*\*\* = $p \leq 0.001$ . While the overall quadratic model appears significant, this may be driven by the linear age component in this model. The first column measuring quadratic $\beta$ coefficients was examined to determine significant quadratic effects which would only be considered meaningful after Bonferroni correction ( $p < .002083$ ).*

**Table V.**
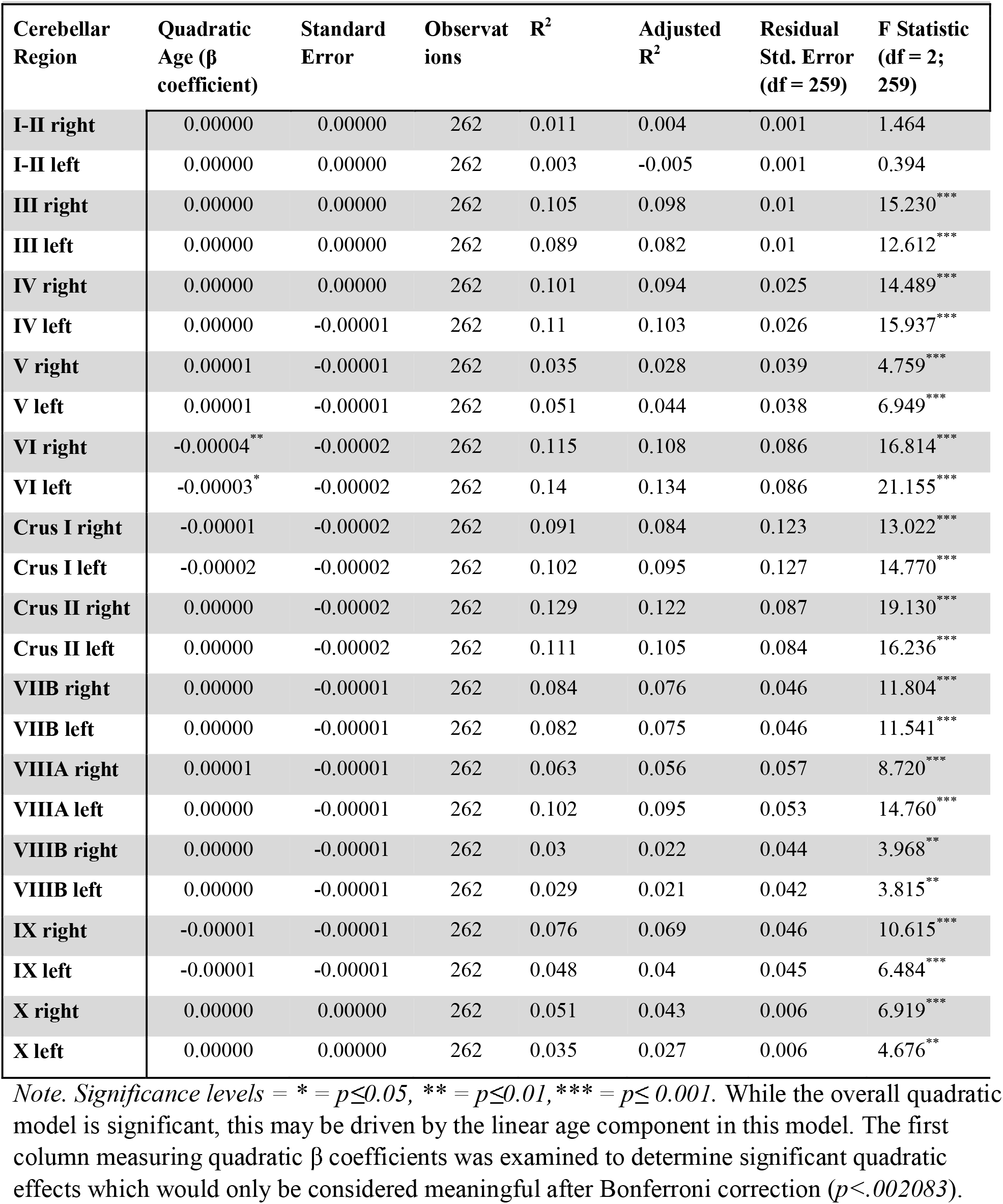
Quadratic Relationship with Age and Adjusted Cerebellar Volume in Females.

Furthermore, comparisons of model fit using AIC did not demonstrate significant differences between linear and quadratic models as specified by a 10-unit difference (Table VI; Bohon & Welch, 2021; Burnham & Anderson, 2004). Although *p*-values and F statistics indicate significance for quadratic models, these values do not indicate that this model is superior to the linear model.

**Table VI.**
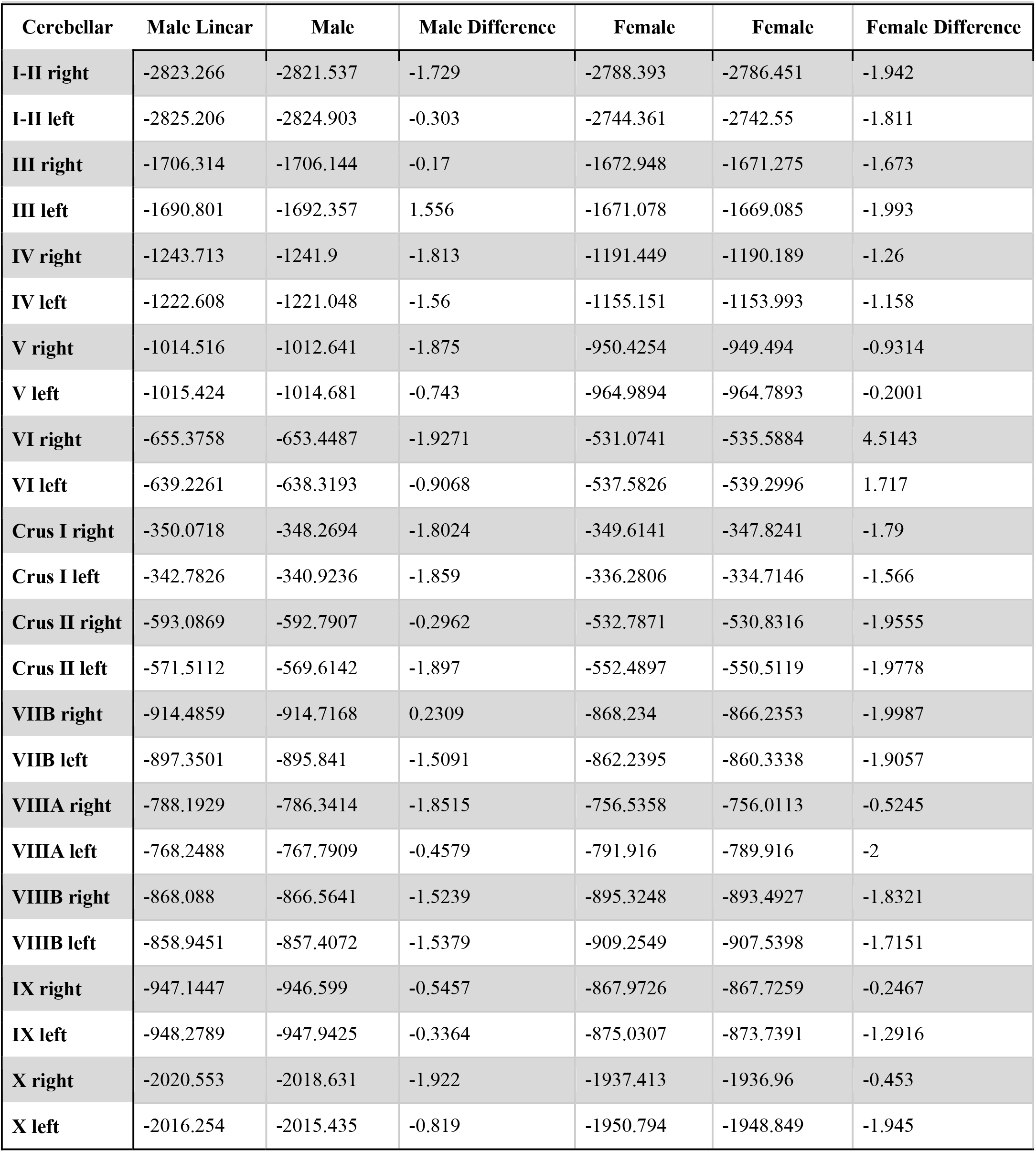
Linear and Quadratic Age Relationships with Adjusted Cerebellar Volume AIC Model Fit Comparisons.

### Reproductive Status

The interactions between sex and reproductive stage investigating their relationship with cerebellar volume were not statistically significant across cerebellar regions [F(3,350) = 0.987, (*p= .483*), please see Figure 3].

**Figure 4.**
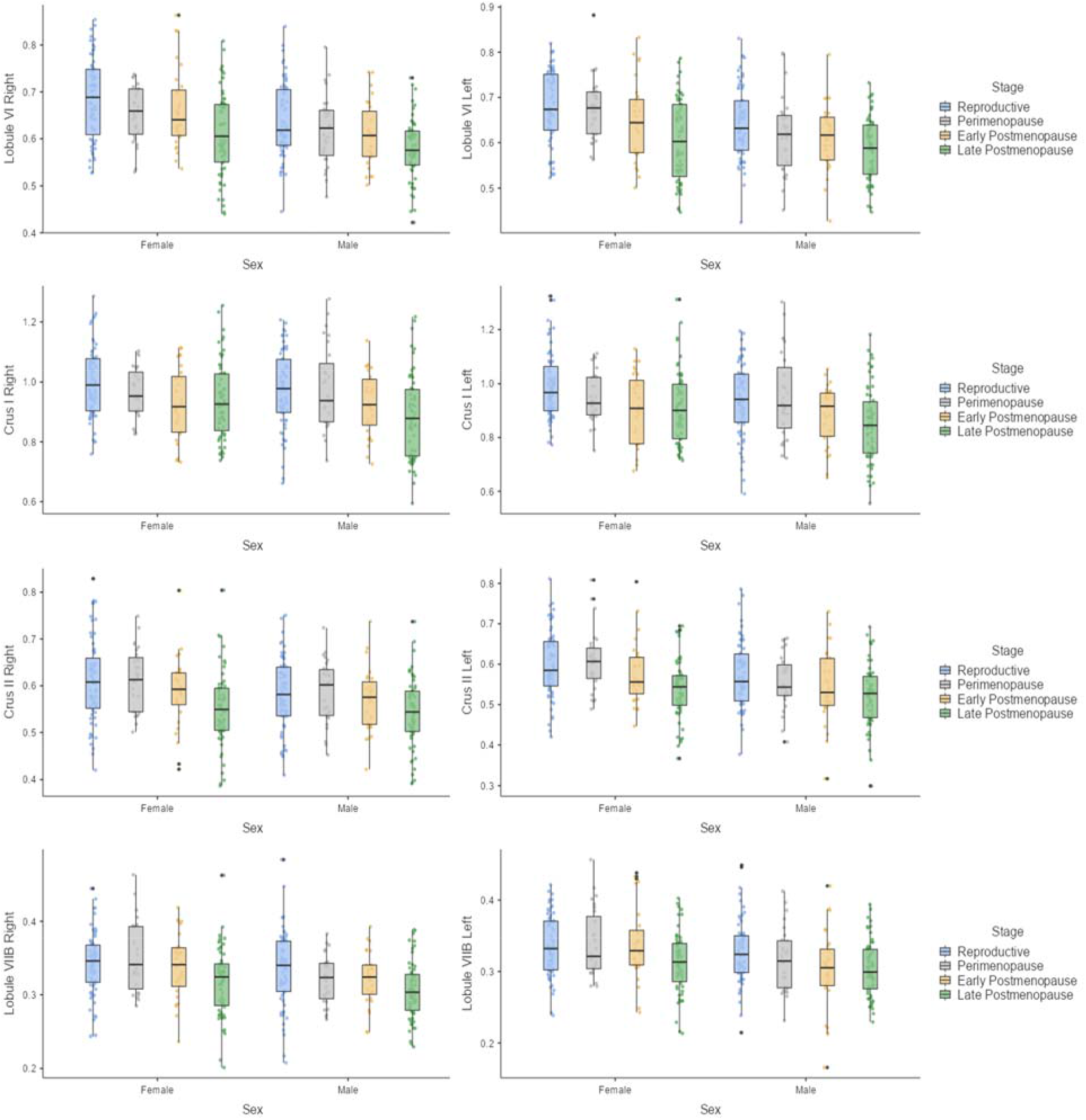
Reproductive Stage Associations with Adjusted Volume in Cerebellar Regions by Sex Mean, interquartile range, and adjusted volume distribution for age-matched males and females in categorized female reproductive stages. There were no significant interactions.

**Figure 5.**
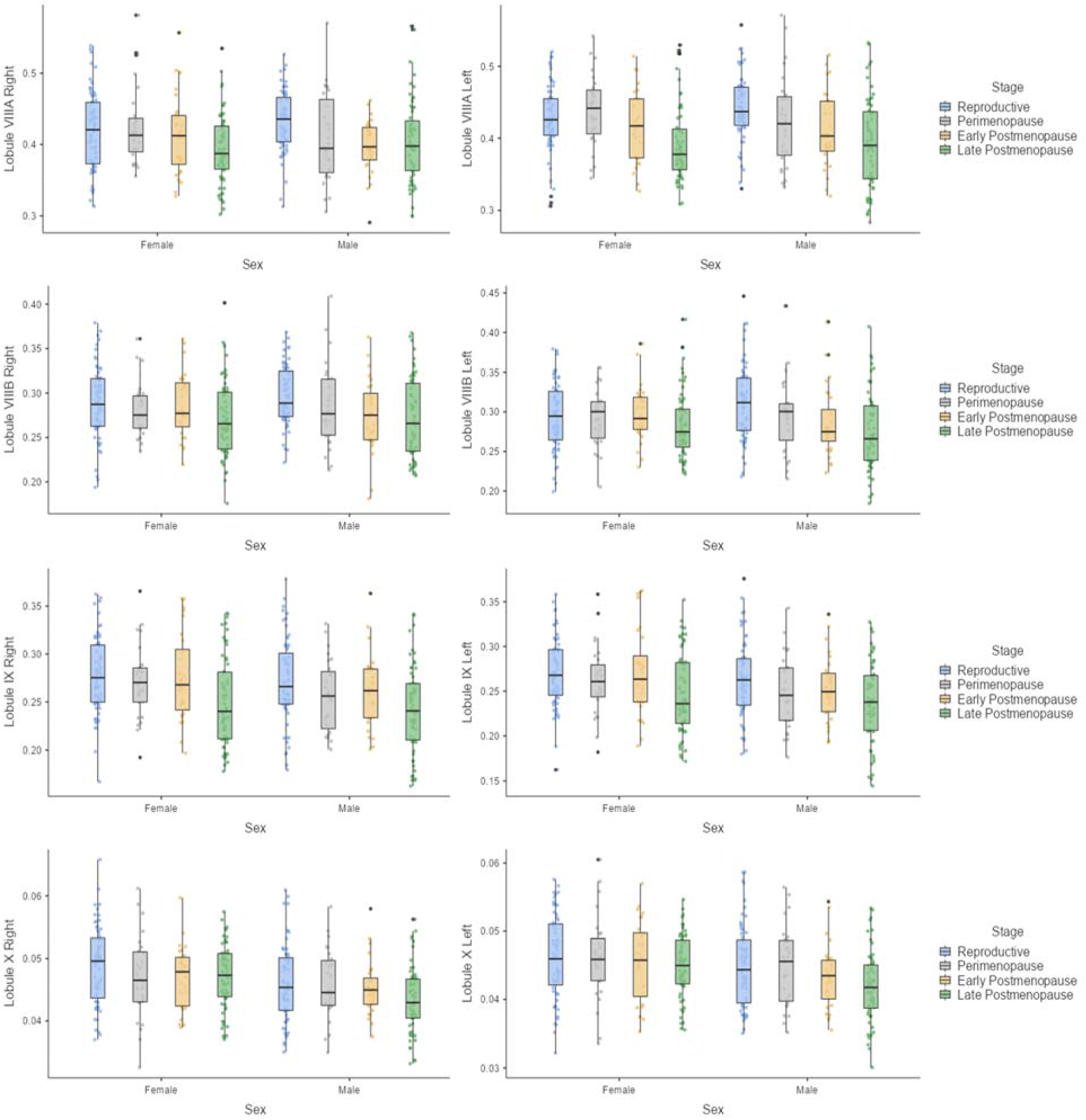
Reproductive Stage Associations with Adjusted Volume in Cerebellar Regions by Sex Mean, interquartile range, and adjusted volume distribution for age-matched males and females in categorized female reproductive stages. There were no significant interactions.

## Discussion

This study investigated regional cerebellar volume to better understand whether age differences and associations with lobular volume differ by sex and as a function of reproductive stage. Specifically, we explored sex differences in regional cerebellar volume as well as differences based on reproductive stage along with linear and quadratic age associations with lobular cerebellar volume in males and females separately. We found significant sex differences as well as significant linear associations across the majority of regions in males and females. However, our results did not suggest superior model fit between linear and quadratic age-volume associations. Lastly, there was no interaction between sex and reproductive stage on regional cerebellar volume. The significant sex differences in regional cerebellar volume--when controlling for age--implies a potential influence of sex hormones on the cerebellum; however, in our assessments based on reproductive stage, that did not come to pass. Significant negative linear relationships between volume and age for each sex in most cerebellar regions could indicate a sex differences in these trajectories over time, although longitudinal data is required to support this theory.

Sex differences in adjusted regional cerebellar volume were found in the majority of lobules examined while controlling for age. These results are largely consistent with past work (Han et al., 2020; Luft et al., 1999; Steele & Chakravaty, 2018) and may suggest that hormones impact cerebellar lobular structure. Consistent with Han and colleagues’ longitudinal sex-related findings in older adults (ages ≥ 50), we found significant sex differences in right Crus II and left Lobule VI (Han et al., 2020). Regarding Han and colleagues’ cross-sectional findings, we did not replicate any areas of significant sex differences (Han et al., 2020). Of note, our study was not an exact replication of this work. We used a different age range and parcellation method (Han et al., 2020). In younger adults (ages 22-36), Steele and Chakravarty reported significant sex differences in several regions including bilateral Crus II (Steele & Chakravaty, 2018), consistent with our work here. Our work provides evidence for structural cerebellar sex differences. Females in our study had greater adjusted cerebellar volume than males in all regions that differed (Figure 1) which was consistent with previous literature (Dimitrova et al., 2006; Weier et al., 2014). Although our data are limited by their cross-sectional nature, this study is the first to examine sex differences in adjusted regional cerebellar volume across the entire adult lifespan in one sample.

While hormonal influences may elucidate nuances in structural cerebellar changes, chronological age has also exhibited an impact on cerebellar volume. The majority of age-volume associations were characterized by significant negative linear relationships with age in both males and females. These findings are consistent with extant literature demonstrating negative linear relationships between cerebellar subregions and age (Han et al., 2020; Koppelmans et al., 2017; Luft et al., 1999; Bernard & Seidler, 2013). However, linear relationships were not entirely consistent between sexes. Right Lobule V, bilateral Lobules VIIIB, and left Lobule X demonstrated significant linear relationships in males which were not seen in females. The sex differences observed here suggest sex-specific aging trajectories may be present, although longitudinal data are required to support this theory. Finally, prior work in adolescent and middle-aged individuals showed quadratic associations with age (Bernard et al., 2015), though we did not see that here after Bonferroni correction. Notably, neither linear nor quadratic models evaluating age-volume associations demonstrated a significantly better fit. Further investigation of cerebellar volume trajectory as it differs between the sexes is warranted in the context of longitudinal studies.

Our initial hypothesis was that regional cerebellar volume would differ based on female reproductive stage given the widely varying hormonal environment. Yet, our study did not reveal any interactions between reproductive stage and sex in the context of cerebellar volume. This was surprising given Mosconi and colleagues’ findings in which females demonstrated significant differences in gray and white matter volumes between reproductive and both perimenopausal and postmenopausal stages (Mosconi et al., 2017). As estrogen therapy has resulted in increased cerebellar gray matter volume in recipients as compared to controls (Boccardi et al., 2006; Ghidoni et al., 2006), the notable declines in estrogen during menopause were expected to have a pronounced influence on cerebellar structure. We speculated that cerebellar regions associated with cognition would be particularly impacted as brain areas associated with cognition are influenced by hormonal fluctuations and cognitive deficits have been associated with hormone levels across reproductive stages (Epperson, Sammel, & Freeman, 2013; Greendale, Derby, & Maki, 2011; Pritschet et al., 2020; Taylor, Pritschet, Yu, & Jacobs, 2019; Rentz et al., 2017; Weber, Maki, & McDermott, 2014). Furthermore, Pritschet and colleagues (2020) demonstrated time synchronous associations between cortical network dynamics and estradiol in a dense-sampling of a single female’s estrous cycle. These changes in the estrous cycle highlight the general impact of hormones on the female brain, and we may in turn speculate that the menopausal transition may also have impacts, given the associated hormonal changes. Examination of the same female over multiple visits (as opposed to a single time point) may be necessary to capture the impact of menopause. That is, the dynamic nature of the resting state signal investigated by Pritschet and colleagues (2020) may be more susceptible to hormonal changes/fluctuations in the short term, while impacts on structure may not be quantifiable until much later after hormone levels have decreased. Furthermore, this study may or may not reflect circulating hormones due to individual variability in the onset of the menopausal transition and the amount sex hormone levels can fluctuate monthly within a female. That is, examination of self-report measures is prone to more variation and inaccuracies than longitudinal hormone assays and structural characterizations of cerebellar regions in each reproductive stage have yet to be established. It is also of course plausible that reproductive stages are unrelated to adjusted regional cerebellar volume, although further investigation is suggested, particularly with longitudinal and hormone data. We speculate that the lack of differences reported here is due in large part to the self-report data used and evaluating a single time point for each participant. Detailed hormonal analyses evaluating each participant at multiple time points across the aging trajectory may be better suited to measure an influence of hormonal environment on the cerebellum.

### Limitations

The Cam-CAN sample has benefits and drawbacks that impact the way the results of this study can be interpreted. One advantage that fewer empirical research studies can boast is that this sample was acquired via population-based recruitment, whereas most samples tend to be overly represented by motivation-based recruitment. However, this was a cross-sectional sample which limits our interpretation of the data, as we cannot investigate change over time. While this dataset is relatively large, evaluating cross-sectional data may not necessarily characterize normal aging processes. As Henson and colleagues described it, the effects of birth-year are not necessarily the same as evaluating true aging within individuals (2020). Furthermore, cohort effects may exist within these data that could have impacted our results.

Our parcellation method differed from previous studies which could make comparisons across studies, more challenging. For instance, MacLullich and colleagues have shown positive correlations between cognitive test performance and segmented vermis volume in older adult males (MacLullich et al., 2004). In our study we examined the vermis as a part of nearby lobular regions, but it was not parceled out. While this region is captured within our study, our results could not be directly compared to previous literature on the vermis.

The STRAW+10 criteria are considered a gold standard in the field of reproductive health and although these criteria exhibit significant strengths, they cannot fully compensate for the limitations of relying solely on self-reported menstrual symptoms in this study (Harlow et al., 2012). The professionals who developed the STRAW+10 criteria recognized that research or clinics may not have access to measurement of endocrine levels or additional confirmatory factors for reproductive stage; thus, the principle criteria only require self-report of menses duration or absence. While the STRAW+10 criteria are valuable to this study, these groupings are ultimately our best approximation based on the self-report data available. Hormonal assays would have provided a more direct and accurate assessment of sex hormone levels improving reproductive stage categorization and allowing for a more direct investigation of relationships between sex steroid hormones and lobular cerebellar volume. Hormonal birth control or treatment of menopausal symptoms also may have confounded our analyses. Ideally, future research would evaluate past and current hormone treatments as well as hormonal assays in the context of longitudinal data collection.

### Conclusions

In summary, this study investigated sex differences in regional cerebellar volume and volume-age associations in addition to the impact of reproductive stage. We found significant sex differences in regional cerebellar volume when controlling for age which suggests a possible influence of sex hormones on the cerebellum. This study also found linear age-volume associations across the majority of cerebellar lobules investigated; however, not all regions showed the same associations across the sexes, which may suggest differential patterns or areas of decline by sex. Aging has been linked to declines in cognitive performance, motor function, and cerebellar volume. Understanding cerebellar volume in the context of sex-differences and healthy aging may provide insight into clinical aging. As incidence of AD is greater in females above and beyond survivor effects, investigation of sex differences as they relate to reproductive stage with longitudinal data may provide greater insight into this pattern. Further investigation into hormonal influences on cerebellar structure and function is warranted for clinical application.

## Acknowledgments

The Cambridge Centre for Ageing and Neuroscience (CamCAN) collected and shared data used in this project. CamCAN funding was provided by the UK Biotechnology and Biological Sciences Research Council (grant number BB/H008217/1), together with support from the UK Medical Research Council and University of Cambridge, UK. This work was further supported by R01AG065010 to J.A.B.

## References

Balsters, J. H., Whelan, C. D., Robertson, I. H., & Ramnani, N. (2013). Cerebellum and cognition: Evidence for the encoding of higher order rules. Cerebral Cortex, 23(6), 1433–1443. https://doi.org/10.1093/cercor/bhs127

Barnes, C., Shechtman, E., Finkelstein, A., & Goldman, D. B. (2009). PatchMatch: A randomized correspondence algorithm for structural image editing. ACM Trans. Graph., 28(3), 24.

Bernard, J. A., Nguyen, A. D., Hausman, H. K., Maldonado, T., Ballard, H. K., Jackson, T. B., … Goen, J. R. (2020). Shaky scaffolding: Age differences in cerebellar activation revealed through activation likelihood estimation meta analysis. Human Brain Mapping, 41(18), 5255–5281.

Bernard, J. A., & Seidler, R. D. (2013). Relationships between regional cerebellar volume and sensorimotor and cognitive function in young and older adults. The Cerebellum, 12(5), 721–737.

Bernard, J. A., & Seidler, R. D. (2013a). Cerebellar contributions to visuomotor adaptation and motor sequence learning: An ALE meta-analysis. Frontiers in Human Neuroscience, 7,27. https://doi.org/10.3389/fnhum.2013.00027

Boccardi, M., Ghidoni, R., Govoni, S., Testa, C., Benussi, L., Bonetti, M., … Frisoni, G. B. (2006). Effects of hormone therapy on brain morphology of healthy postmenopausal women: a Voxel-based morphometry study. Menopause, 13(4), 584–591.

Bohon, C., & Welch, H. (2021). Quadratic relations of BMI with depression and brain volume in children: Analysis of data from the ABCD study. Journal of Psychiatric Research, 136, 421–427.

Boyle, C. P., Raji, C. A., Erickson, K. I., Lopez, O. L., Becker, J. T., Gach, H. M., … Thompson, P. M. (2021). Estrogen, brain structure, and cognition in postmenopausal women. Human brain mapping, 42(1), 24–35.

Buckler, H. (2005). The menopause transition: endocrine changes and clinical symptoms. British Menopause Society Journal, 11(2), 61–65.

Buckner, R. L., Head, D., Parker, J., Fotenos, A. F., Marcus, D., Morris, J. C., & Snyder, A. Z. (2004). A unified approach for morphometric and functional data analysis in young, old, and demented adults using automated atlas-based head size normalization: reliability and validation against manual measurement of total intracranial volume. Neuroimage, 23(2), 724–738.

Burnham, K. P., & Anderson, D. R. (2004). Multimodel inference: understanding AIC and BIC in model selection. Sociological methods & research, 33(2), 261–304.

Carass, A., Cuzzocreo, J. L., Han, S., Hernandez-Castillo, C. R., Rasser, P. E., Ganz, M., … Thyreau, B. (2018). Comparing fully automated state-of-the-art cerebellum parcellation from magnetic resonance images. Neuroimage, 183, 150–172.

Chen, S. H. A., & Desmond, J. E. (2005). Cerebrocerebellar networks during articulatory rehearsal and verbal working memory tasks. NeuroImage, 24(2), 332–338. https://doi.org/10.1016/j.neuroimage.2004.08.032

de Dieu Uwisengeyimana, J., Nguchu, B. A., Wang, Y., Zhang, D., Liu, Y., Qiu, B., & Wang, X. (2020). Cognitive function and cerebellar morphometric changes relate to abnormal intra-cerebellar and cerebro-cerebellum functional connectivity in old adults. Experimental Gerontology, 140, 111060.

Dimitrova, A., Zeljko, D., Schwarze, F., Maschke, M., Gerwig, M., Frings, M., … Timmann, D. (2006). Probabilistic 3D MRI atlas of the human cerebellar dentate/interposed nuclei. Neuroimage, 30(1), 12–25.

Epperson, C. N., Sammel, M. D., & Freeman, E. W. (2013). Menopause effects on verbal memory: findings from a longitudinal community cohort. The Journal of clinical endocrinology & metabolism, 98(9), 3829–3838.

Ghidoni, R., Boccardi, M., Benussi, L., Testa, C., Villa, A., Pievani, M., … Binetti, G. (2006). Effects of estrogens on cognition and brain morphology: involvement of the cerebellum. Maturitas, 54(3), 222–228.

Greendale, G. A., Derby, C. A., & Maki, P. M. (2011). Perimenopause and cognition. Obstetrics and Gynecology Clinics, 38(3), 519–535.

Harada, C. N., Love, M. C. N., & Triebel, K. L. (2013). Normal cognitive aging. Clinics in geriatric medicine, 29(4), 737–752.

Harlow, S. D., Gass, M., Hall, J. E., Lobo, R., Maki, P., Rebar, R. W., … STRAW+ 10 Collaborative Group. (2012).Executive summary of the Stages of Reproductive Aging Workshop+ 10: addressing the unfinished agenda of staging reproductive aging. The Journal of Clinical Endocrinology & Metabolism, 97(4), 1159–1168.

Hedges, V. L., Ebner, T. J., Meisel, R. L., & Mermelstein, P. G. (2012). The cerebellum as a target for estrogen action. Frontiers in neuroendocrinology, 33(4), 403–411.

Hlavac, M. (2018). stargazer: Well-Formatted Regression and Summary Statistics Tables. Central European Labour Studies Institute (CELSI). https://CRAN.R-project.org/package=stargazer

Hulst, T., van der Geest, J. N., Thürling, M., Goericke, S., Frens, M. A., Timmann, D., & Donchin, O. (2015). Ageing shows a pattern of cerebellar degeneration analogous, but not equal, to that in patients suffering from cerebellar degenerative disease. Neuroimage, 116, 196–206.

Jernigan, T. L., Archibald, S. L., Fennema-Notestine, C., Gamst, A. C., Stout, J. C., Bonner, J., & Hesselink, J. R. (2001). Effects of age on tissues and regions of the cerebrum and cerebellum. Neurobiology of aging, 22(4), 581–594.

Kikkert, L. H., Vuillerme, N., van Campen, J. P., Hortobágyi, T., & Lamoth, C. J. (2016). Walking ability to predict future cognitive decline in old adults: a scoping review. Ageing research reviews, 27, 1–14.

King, M., Hernandez-Castillo, C. R., Poldrack, R. A., Ivry, R. B., & Diedrichsen, J. (2019). Functional boundaries in the human cerebellum revealed by a multi-domain task battery. Nature neuroscience, 22(8), 1371–1378.

Kluger, A., Gianutsos, J. G., Golomb, J., Ferris, S. H., George, A. E., Franssen, E., & Reisberg, B. (1997). Patterns of motor impairment in normal aging, mild cognitive decline, and early Alzheimer’Disease. The Journals of Gerontology Series B: Psychological Sciences and Social Sciences, 52(1), P28–P39.

Koppelmans, V., Hoogendam, Y. Y., Hirsiger, S., Mérillat, S., Jäncke, L., & Seidler, R. D. (2017). Regional cerebellar volumetric correlates of manual motor and cognitive function. Brain Structure and Function, 222(4), 1929–1944.

Lezak, M. D., Howieson, D. B., Loring, D. W., & Fischer, J. S. (2004). Neuropsychological assessment. Oxford University Press, USA.

Luft, Andreas R., et al. “Patterns of age-related shrinkage in cerebellum and brainstem observed in vivo using three-dimensional MRI volumetry.” Cerebral Cortex 9.7 (1999): 712–721.

MacLullich, A. M., Edmond, C. L., Ferguson, K. J., Wardlaw, J. M., Starr, J. M., Seckl, J. R., & Deary, I. J. (2004). Size of the neocerebellar vermis is associated with cognition in healthy elderly men. Brain and cognition, 56(3), 344–348.

Marquis, S., Moore, M. M., Howieson, D. B., Sexton, G., Payami, H., Kaye, J. A., & Camicioli, R. (2002). Independent predictors of cognitive decline in healthy elderly persons. Archives of neurology, 59(4), 601–606.

Miller, T. D., Ferguson, K. J., Reid, L. M., Wardlaw, J. M., Starr, J. M., Seckl, J. R., … MacLullich, A. M. (2013). Cerebellar vermis size and cognitive ability in community-dwelling elderly men. The Cerebellum, 12(1), 68–73.

Morrison, J. H., & Baxter, M. G. (2012). The ageing cortical synapse: hallmarks and implications for cognitive decline. Nature Reviews Neuroscience, 13(4), 240–250.

Morrison, J. H., Brinton, R. D., Schmidt, P. J., & Gore, A. C. (2006). Estrogen, menopause, and the aging brain: how basic neuroscience can inform hormone therapy in women. Journal of Neuroscience, 26(41), 10332–10348.

Mosconi, L., Berti, V., Guyara-Quinn, C., McHugh, P., Petrongolo, G., Osorio, R. S., … Brinton, R. D. (2017). Perimenopause and emergence of an Alzheimer’s bioenergetic phenotype in brain and periphery. PloS one, 12(10), e0185926.

Munro, C. A., Winicki, J. M., Schretlen, D. J., Gower, E. W., Turano, K. A., Muñoz, B., … West, S. K. (2012). Sex differences in cognition in healthy elderly individuals. Aging, Neuropsychology, and Cognition, 19(6), 759–768.

Pillerová, M., Borbélyová, V., Hodosy, J., Riljak, V., Renczés, E., Frick, K. M., & Tóthová, L’. (2021). On the role of sex steroids in biological functions by classical and non-classical pathways. An update. Frontiers in Neuroendocrinology, 100926.

Pritschet, L., Santander, T., Taylor, C. M., Layher, E., Yu, S., Miller, M. B., … Jacobs, E. G. (2020). Functional reorganization of brain networks across the human menstrual cycle. NeuroImage, 220, 117091.

Raz, N., Dupuis, J. H., Briggs, S. D., McGavran, C., & Acker, J. D. (1998). Differential effects of age and sex on the cerebellar hemispheres and the vermis: a prospective MR study. American Journal of Neuroradiology, 19(1), 65–71.

Raz, N., Ghisletta, P., Rodrigue, K. M., Kennedy, K. M., & Lindenberger, U. (2010). Trajectories of brain aging in middle-aged and older adults: regional and individual differences. Neuroimage, 51(2), 501–511.

Raz, N., Gunning-Dixon, F., Head, D., Williamson, A., & Acker, J. D. (2001). Age and sex differences in the cerebellum and the ventral pons: a prospective MR study of healthy adults. American Journal of Neuroradiology, 22(6), 1161–1167.

R Core Team (2021). R: A language and environment for statistical computing. R Foundation for Statistical Computing, Vienna, Austria. URL https://www.R-project.org/.

Rentz, D. M., Weiss, B. K., Jacobs, E. G., Cherkerzian, S., Klibanski, A., Remington, A., … Goldstein, J. M. (2017). Sex differences in episodic memory in early midlife: impact of reproductive aging. Menopause (New York, NY), 24(4), 400.

Rezaee, Z., & Dutta, A. (2020). Lobule specific dosage considerations for cerebellar transcranial direct current stimulation during healthy aging: A computational modeling study using age specific magnetic resonance imaging templates. Neuromodulation: Technology at the Neural Interface, 23(3), 341–365.

Robertson, D., Craig, M., Van Amelsvoort, T., Daly, E., Moore, C., Simmons, A., … Murphy, D. (2009). Effects of estrogen therapy on age-related differences in gray matter concentration. Climacteric, 12(4), 301–309.

Romero, J. E., Coupé, P., Giraud, R., Ta, V. T., Fonov, V., Park, M. T. M., … Manjón, J. V. (2017). CERES: a new cerebellum lobule segmentation method. Neuroimage, 147, 916–924.

Sakamoto, Y., Ishiguro, M., and Kitagawa G. (1986). Akaike Information Criterion Statistics. D. Reidel Publishing Company.

Savica, R., Wennberg, A., Hagen, C., Edwards, K., Roberts, R. O., Hollman, J. H., … Mielke, M. M. (2017). Comparison of gait parameters for predicting cognitive decline: the Mayo Clinic Study of Aging. Journal of Alzheimer’s Disease, 55(2), 559–567.

Schaefer, S., & Schumacher, V. (2011). The interplay between cognitive and motor functioning in healthy older adults: findings from dual-task studies and suggestions for intervention. Gerontology, 57(3), 239–246.

Seidler, R. D., Bernard, J. A., Burutolu, T. B., Fling, B. W., Gordon, M. T., Gwin, J. T., … Lipps, D. B. (2010). Motor control and aging: links to age-related brain structural, functional, and biochemical effects. Neuroscience & Biobehavioral Reviews, 34(5), 721–733.

Shafto, M. A., Tyler, L. K., Dixon, M., Taylor, J. R., Rowe, J. B., Cusack, R., … Henson, R. N. (2014). The Cambridge Centre for Ageing and Neuroscience (Cam-CAN) study protocol: a cross-sectional, lifespan, multidisciplinary examination of healthy cognitive ageing. BMC neurology, 14(1), 204.

Stoodley, C. J., Valera, E. M., & Schmahmann, J. D. (2012). Functional topography of the cerebellum for motor and cognitive tasks: An fMRI study. NeuroImage, 59, 1560–1570. https://doi.org/10.1016/j.neuroimage.2011.08.065

Stoodley, C. J., & Schmahmann, J. D. (2009). Functional topography in the human cerebellum: A meta-analysis of neuroimaging studies. NeuroImage, 44(2), 489–501. https://doi.org/10.1016/j.neuroimage.2008.08.039

Ta, V. T., Giraud, R., Collins, D. L., & Coupé, P. (2014, September). Optimized patchmatch for near real time and accurate label fusion. In International Conference on Medical Image Computing and Computer-Assisted Intervention (pp. 105–112). Springer, Cham.

Taylor, C. M., Pritschet, L., Yu, S., & Jacobs, E. G. (2019). Applying a women’s health lens to the study of the aging brain. Frontiers in human neuroscience, 13, 224.

Taylor, J. R., Williams, N., Cusack, R., Auer, T., Shafto, M. A., Dixon, M., … Henson, R. N. (2017). The Cambridge Centre for Ageing and Neuroscience (Cam-CAN) data repository: Structural and functional MRI, MEG, and cognitive data from a crosssectional adult lifespan sample. Neuroimage, 144, 262–269.

The jamovi project (2021). jamovi (Version 1.6) [Computer Software]. Retrieved from https://www.jamovi.org

Vest, R. S., & Pike, C. J. (2013). Gender, sex steroid hormones, and Alzheimer’s disease. Hormones and behavior, 63(2), 301–307.

Weber, M. T., Maki, P. M., & McDermott, M. P. (2014). Cognition and mood in perimenopause: a systematic review and meta-analysis. The Journal of steroid biochemistry and molecular biology, 142, 90–98.

Weier, K., Fonov, V., Lavoie, K., Doyon, J., & Collins, D. L. (2014). Rapid automatic segmentation of the human cerebellum and its lobules (RASCAL)—Implementation and application of the patch▫based▫labed fusion technique with a template library to segment the human cerebellum. Human brain mapping, 35(10), 5026–5039.

Wickham, H. (2016). Ggplot2: Elegant graphics for data analysis (2nd ed.) [PDF]. Springer International Publishing.

Woodruff-Pak, D. S., Foy, M. R., Akopian, G. G., Lee, K. H., Zach, J., Nguyen, K. P. T., … Thompson, R. F. (2010). Differential effects and rates of normal aging in cerebellum and hippocampus. Proceedings of the National Academy of Sciences, 107(4), 1624–1629.

Xu, J., Kobayashi, S., Yamaguchi, S., Iijima, K. I., Okada, K., & Yamashita, K. (2000). Gender effects on age-related changes in brain structure. American journal of neuroradiology, 21(1), 112–118.

